# Insect abundance and richness response to ecological reclamation on well pads 5-12 years into succession in a semi-arid natural gas field

**DOI:** 10.1101/2024.05.03.592291

**Authors:** Michael F. Curran, Jasmine Allison, Timothy J. Robinson, Blair L. Robertson, Alexander H. Knudson, Bee M.M. Bott, Steven Bower, Bobby M. Saleh

## Abstract

Natural gas extraction is a critical driver of the economy in western North America. Ecological reclamation is important to ensure surface disturbance impacts associated with natural gas development are not permanent and to assist native biota. Previous studies in semi-arid natural gas fields within Sublette County, Wyoming, USA have shown insects respond favorably to 1-3-year-old well pads undergoing reclamation compared to older successional, reference vegetation communities dominated by Wyoming big sagebrush (*Artemisia tridenta* spp. *Wyomingensis*). Here we examined well pads which were initially seeded between 5, 8, 10, 11, and 12 prior to our study. We used a free, image-based software called SamplePoint to quantify vegetation on these well pads and adjacent reference areas from high-resolution cell phone camera pictures. Insects were collected with a sweep net and identified to the family and morphospecies level. Statistical analyses were conducted to compare both vegetation and insect communities between reclamation sites and their paired reference area. We found little statistical difference between vegetation communities across our study, but found significantly more insect abundance in 3 of 5 years and significantly higher family and species richness in 4 of 5 years. Our results suggest reclamation of natural gas well pads within an old successional stand of sagebrush continues to support higher levels of insect biodiversity and abundance for at least 12 years. As insects are the most diverse group of animals on Earth and because they provide a wide array of ecosystem services, our findings suggest ecological reclamation plays an important role in returning biodiversity and ecosystem functionality to a semi-arid and old successional sagebrush-steppe ecosystem.

## 1. Introduction

Ecological reclamation is a necessary process to aid native biota after land development requiring surface disturbance has taken place [1]. Although drylands make up ~40% of the Earth’s surface [2] and are home to ~39% of the human population [3], research from drylands only accounts for <6% of published terrestrial ecology studies [4] and <5% of published studies related to restoration ecology [5]. The expanse of drylands is predicted to increase with changing climate conditions and a growing human population will continue to rely on natural resources for the foreseeable future [6, 7]. Wyoming, the 9th largest state in the United States by land mass, has climates ranging from arid to semi-arid across its vast expanse [8]. Although it is the least populated state, Wyoming is rich in natural resources and home to a wide array of biodiversity and wildlife [8]. Since biodiversity loss is known to have negative impacts on humanity and because natural resource extraction is often linked to land surface disturbance [8, 9], understanding how biodiversity responds to land surface disturbance and reclamation activities associated with resource extraction is paramount.

Natural gas provides tangible benefits to humans and production of the resource is a major driver of Wyoming’s economy [8]. In order to extract natural gas, well pads are typically constructed by stripping and stockpiling soil and removing vegetation from an area to provide a safe working environment and to house equipment necessary to build a well [10]. After well construction is complete, the construction area is typically decompacted before soil from stockpiles is respread across the non-active area (~75%) of the well pad and then seeded with native vegetation to begin the process of interim reclamation [10, 11]. In the Pinedale Anticline natural gas field (Sublette County, Wyoming, USA), operators are required to comply with reclamation regulatory criteria, which vary between and among State and Federal Government agencies [12, 13]. These criteria are focused on erosion control, minimizing invasive and noxious vegetation species, and establishing native vegetation communities which are similar to adjacent reference ecosystems which have not been directly impacted by natural gas activities [11-13]. In the past decade, efforts to exceed minimum regulatory criteria efforts towards restoring wildlife habitat are common practice in Wyoming [8].

Efforts to reclaim dryland ecosystems are challenged by a variety of environmental factors including invasive vegetation [14], low and unpredictable precipitation [8], lack of available seed resources [15], poor soils [16], and extreme temperatures [17]. Situated in a sagebrush-steppe ecosystem with elevations above 2,100m in an area receiving 254-356 mm mean annual precipitation (MAP) and experiencing 42 frost-free days per year, many of these challenges are exacerbated in the Pinedale Anticline natural gas field. Aside from environmental and climatic challenges, operators in the area must deal with multiple land use goals (e.g., domestic livestock grazing, recreation, wildlife habitat) and abide by aforementioned regulatory criteria. Aside from erosion control and lack of noxious or invasive vegetation, operators in the Pinedale Anticline are required to achieve >50% of shrub density compared to an adjacent reference area. Typically, reference sites in the area are comprised of old stands of Wyoming big sagebrush (*Artemisia tridentata* spp. *Wyomingensis*) which have been subject to domestic livestock grazing since the 1870s [18]. Research has shown sagebrush establishment in arid and semiarid environments may take decades [19]. As soil alterations associated with natural gas well pad development result in a pulse of plant-available nitrogen which sagebrush are not well adapted to [20, 21], operators in the area have achieved better results combating annual invasive weed species and stabilizing locations using diverse seed mixes comprised of native annual forb and perennial forb, grass and shrub species [22]. This can be looked at as a form of ‘assisted succession’ [23], setting a pathway for the eventual establishment of mature sagebrush. While careful reclamation planning, soil management, and seed mix application are required for initial vegetation establishment in arid environments [17, 20], establishing plant-pollinator networks is critical to ensure these vegetation communities are self-sustaining over time [24-27].

While some plants in rangeland ecosystems rely mainly on wind pollination [28], over 75% of flowering plants rely on animal pollination to successfully reproduce and maintain genetic diversity [29]. Aside from their critical role as pollinators, insects are the most abundant and diverse group of animals on Earth and provide numerous important ecosystem services [30-33]. In addition to pollination and biodiversity, other ecosystem services insects provide include nutrient cycling and decomposition [34, 35], nutrition of higher food levels [36], and pest control [37]. Despite their importance, terrestrial insects have been declining globally [38] and are often overlooked in terrestrial wildlife studies [39]. Recent research suggests reductions in invertebrates are associated with declining ecosystem performance [40].

Giving cause for hope, a 2018 meta-analysis suggests habitat restoration efforts are beneficial to wild pollinators [41]. Measuring insect richness and diversity has shown to be a sound indicator of ecosystem functionality and restoration success since insects respond rapidly to environmental change, provide more ecosystem services than other animals, and because statistically valid samples can be obtained in a short time period [42-43]. Efforts to understand insect response to restoration efforts are heavily focused on pollinators in crop agriculture ecosystems [44] and very few studies have been conducted to determine how insects respond to reclamation associated with oil and natural gas development [45-47]. Previous studies examining how insects respond to reclamation practices on Sublette County natural gas well pads show well pads exhibiting early successional vegetation (1-3 years post reclamation initiation) contain higher insect abundance and diversity than adjacent reference ecosystems [46, 47]. To date, no studies have examined how insect abundance and diversity fare on reclaimed well pads with vegetation communities in later successional stages.

The objective of this study is to examine differences between vegetation communities on natural gas well pads on which reclamation was initiated 5-12 years ago compared to adjacent reference communities as well as to examine differences in insect communities on those sites and their paired reference sites. A total of 15 natural gas well pads (3 each in which reclamation was initiated 5, 8, 10, 11, and 12 years ago) were examined along with a paired reference site. All reference areas are dominated by old stands of Wyoming big sagebrush with understories containing few grass and flowering species, whereas reclaimed well pad locations contained more diverse vegetation communities. Based on previous research examining vegetation and insects in this area [46, 47], we hypothesized that vegetation communities on reclaimed well pads would be different than reference sites and that insect abundance and diversity would be higher on reclaimed well pads than reference sites. As information about insects in the area is limited, we also sought to determine if different insect families were distributed equally among reclaimed sites and reference areas.

## 2. Materials and Methods

### 2.1 Study Area

The Pinedale Anticline natural gas field is located near the town of Pinedale in Sublette County, WY, USA (Figure 1). The entire gas field is located above 2100m and is considered semi-arid, with a MAP of 254-365mm. The area has an average of 42 frost-free days per year and experiences highly variable daily temperatures during the short growing season, with daily minimum temperatures ranging between -1 and 6 ° C and daily maximum temperatures ranging between 19 and 26 ° C. Despite the challenging environment, the Pinedale Anticline is home to wildlife including Greater sage-grouse (*Centrocercus urophasianus*), mule deer (*Odocoileus hemionus*), pronghorn antelope (*Antilocapra americana*), and a wide variety of songbirds and other sagebrush obligate species.

**Figure 1.**
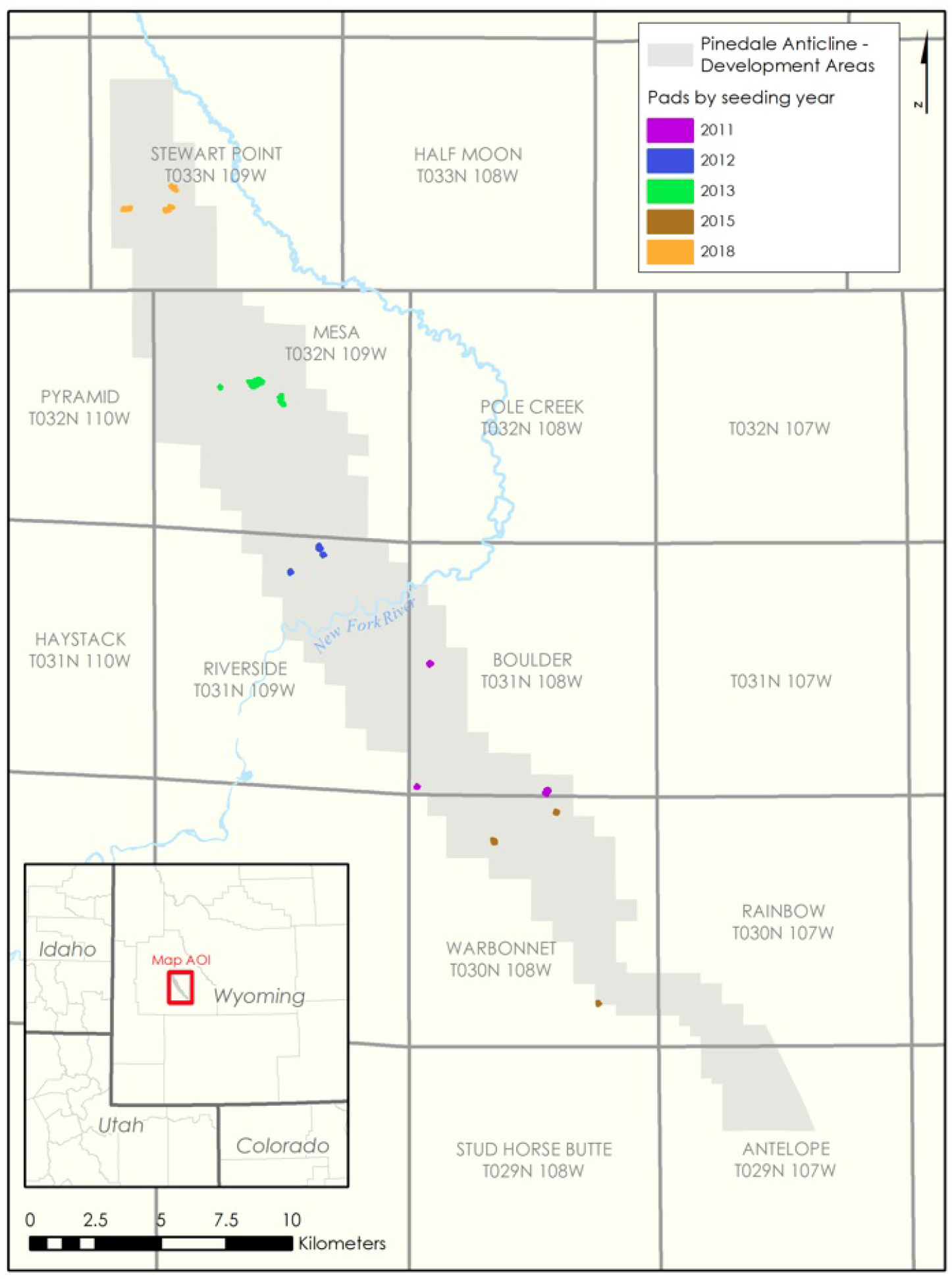
A map depicting the area of the Pinedale Anticline natural gas field with study locations color coded by seeding year.

Natural gas development began in the Pinedale Anticline ca. 2000, with predominantly 5 different upstream (i.e., exploration and production) operating companies managing different sections of the gas field. Reclamation regulatory criteria are set in place by the Wyoming Department of Environmental Quality (WDEQ) and the DOI BLM 2008 Record of Decision (ROD) for Supplemental Environmental Impact Statement (SEIS) Pinedale Anticline Project Area (PAPA). The BLM 2008 ROD SEIS PAPA created the Pinedale Anticline Project Office (PAPO) to oversee and manage mitigation and monitoring and the funding for mitigation and monitoring. The PAPO is made up of interagency governmental consortium consisting of the WDEQ, Wyoming Game and Fish Department, Wyoming Department of Agriculture, Sublette County Conservation District, and the Bureau of Land Management. The PAPO oversees Pinedale Anticline Data Management System (PADMS) which is a reclamation data monitoring tool. The WDEQ Stormwater Pollution Prevention Plan requires reclaimed well pad sites to be stabilized as evidenced by 70% vegetation cover compared to an adjacent, undisturbed reference system as well as a lack of erosion features present in the reclamation area. The 2008 ROD SEIS PAPA reclamation regulatory criteria suggest reclaimed well pad locations must (1) be in stable condition with no erosion features, (2) have a minimum of 3 native perennial grass species of which at least 2 are bunchgrasses, (3) be free from all species listed as noxious in WY, (4) have a resilient vegetation community as evidenced by well-developed root systems, flowers and seed heads, (5) and meet several criteria based on comparison to an adjacent, undisturbed reference area. These criteria include that the reclamation site during interim reclamation which is the stage prior to final reclamation, must have the following in comparison to a reference site: (1) an average density or frequency of forbs which is >75% within 5 years, (2) an average density of frequency of shrubs which is >50% within 5 years, and (3) a percentage of bareground cover less than or equal. While all operators in the area are subject to the same criteria, initial reclamation plans and practices were unique to different operators. More recently, the natural gas well pad locations in the Pinedale Anticline have been acquired by one operating company who actively manages all locations.

Soils throughout the Pinedale Anticline are predominantly sandy loams. While reference sites are unique to each well pad location, the majority of reference sites throughout the field are dominated by old stands of Wyoming big sagebrush with a mixed grass understory which matches the United States Department of Agriculture – Natural Resources Conservation Service’s ecological site description characterization of the historic plant climax community. According to definitions by the Society for Ecological Restoration, reference areas in the Pinedale Anticline natural gas field can be considered cultural reference ecosystems [48] since they have been managed and grazed by domestic livestock long before natural gas development began in the area [18].

### 2.2 Site Selection

The Pinedale Anticline natural gas field was stratified by management area to assist with site selection (Figure 1). This gas field was developed by different operators and development was not consistent between 2000-2023 (e.g., some years saw more well pads constructed than others). Three sites were randomly selected from five management units after each management unit was grouped by year of initial reclamation and seed mix used in effort to obtain replicates based on similar vegetation communities at the time of sampling (Figure 1). Sites in the Boulder management area were seeded in 2011, sites in the Riverside management area were seeded in 2012, sites in the Mesa management area were seeded in 2013, sites in the Warbonnet management area were seeded in 2015, and sites in the Stewart Point management area were seeded in 2018. While seed mixes varied across management units, they were consistent within management units and all species within each seed mix are native to Sublette County, WY and approved by the BLM with goals of supporting multiple land uses for wildlife and domestic livestock (seed mixes unique to each area are available in Appendix A). The locations selected within the Boulder management area ranged from 2125-2197 m elevation, whereas sites in the Riverside management area ranged from 2137-2150 m elevation, sites in the Mesa management area ranged from 2273-2277 m elevation, sites in the Warbonnet management area ranged from 2163-2231 m elevation, and sites in the Stewart Point management area ranged from 2204-2356 m elevation.

Vegetation and insect sampling were conducted concurrently during peak flowering between 14-16 August 2023. All sampling was conducted by author M.F.C. in sunshine conditions with wind speeds <6 kph between 0900-1200. Temperatures during sampling events ranged between 13-21 o C on 14 August, between 13-22 o C on 15 August and between 14-23 o C on 16 August.

### 2.3 Vegetation Sampling

A matched pair study design was used in this effort with each well pad and its paired reference site having two 40 m transects sampled at 10 m and 20 m away from the well pad edge. This design is similar to previous research [46, 47] and is in accordance with the 2008 ROD SEIS PAPA. Along each transect 0.5 m2 images were taken at 5 m intervals, resulting in 9 images per transect or 18 images per each well pad and paired reference location. All images were taken by hand with a 50 mega-pixel camera on a Samsung Galaxy A14 taking images perpendicular to the ground (freehand, nadir technique) [49]. The ground sample distance in each photo was ~0.2 mm, which is similar to previously published research using image analysis to identify vegetation on reclamation sites and in rangeland areas [13, 50-51]. A 5×5 grid was placed inside of each photo using a free image analysis software called ‘SamplePoint’ [52] to examine 25 points per photo as recommended by Ancin-Murguzur et al. [53], resulting in 225 data points per transect or 450 data points for each well pad and paired reference area. Each data point was classified as bareground (i.e., soil or rock), litter, or vegetation to the species-specific level. Vegetation at the species-specific level was classified as native or non-native and then grouped by life form (i.e., grass, forb, or shrub) prior to statistical analysis.

### 2.4 Insect Sampling

Insect sampling was conducted along the same transects during the same days and times as vegetation sampling. Forty sweeps were taken along each 40 m transect location using a canvas sweep net with a 38 cm diameter and 50 cm handle. These sweeps were conducted prior to vegetation sampling to minimize potential disruption to insects from observer being on site. After each transect sweep was completed, insects were placed in a Zip-Lock ® bag and placed in a cooler containing dry ice which was a minimum of 200 m from the sampling locations. A minimum of a 10-minutes was spent at this distance before walking back to the sampling area to complete another sweep net sample to minimize any ‘flushing’ effects on insects [54].

After insect sampling was complete, all insects were shipped in coolers to the Utah State University Plant Pest Diagnostics Laboratory in Logan, Utah, United States. Author A.H.K., who has a PhD in Entomology, identified insects to the family level and organized each family into morphospecies. The majority of insects were identified to family using Triplehorn & Johnson [55], with the Diptera identified to family using McAlpine et. al [56]. Diverse superfamilies (i.e., *Chalcidoidea* and *Gellichoidea*) were kept to superfamily save the nominal families. Insect species were then assigned individual morphospecies numbers and individuals of each morphospecies were counted at each site. Immature insects were associated with corresponding adult specimens except Lepidoptera and are included in the count totals for each morphospecies per site. Voucher specimens have been deposited to the University of Wyoming Insect Museum.

### 2.5 Statistical Analysis

As each management area varied in elevation and proximity to the river in the southern portion of the Pinedale Anticline natural gas field (Figure 1) and because insect sampling may be heavily influenced by a number of factors aside from vegetation (e.g., elevation, proximity to other resources, time of day, temperature, etc.) [57-59], we did not attempt to correlate insect abundance and diversity with vegetation cover. Instead, vegetation analysis and insect analysis were conducted independently among study areas described in section 2.2.

Vegetation analysis was conducted using automatically generated .csv files from SamplePoint. Pairwise t-comparisons with Bonferroni adjustments, were used to compare each cover class category across reclaimed and reference sites for each of the five locations. The Bonferroni adjusted p-values are provided in Table 1.

**Table 1.** Paired t-comparisons with Bonferroni adjustments of reclaimed vs. reference at each site (*Note that there was no recorded ‘non-native’ observations for Stewart Point)

| Cover Type | Boulder | Riverside | Mesa | Warbonnet | Stewart Point |
| --- | --- | --- | --- | --- | --- |
| **Bareground** | 0.201 | **0.028** | **0.025** | 0.063 | 0.575 |
| **Forb** | 0.130 | 0.253 | 0.291 | 0.202 | 0.057 |
| **Grass** | 0.109 | 0.211 | 0.555 | 0.324 | **0.004** |
| **Litter** | 0.109 | 0.483 | 0.505 | 0.375 | 0.850 |
| **Non-Native** | 0.704 | 0.270 | 0.423 | 0.742 | NA |
| **Shrub** | 0.223 | **0.047** | 0.603 | 0.065 | 0.104 |

For insect data, generalized linear model regressions were run to determine potential relationships between the response (insect abundance, species richness, and family richness, respectively) and site location, reclamation status and the interaction of site location and reclamation status. Poisson models were run for species richness and family richness and due to overdispersion, a negative binomial regression was utilized for modeling insect abundance. Parametric bootstrapping was then implemented to provide 95% confidence intervals on the predicted mean response for each combination of site type and reclamation status. Additionally, the top three insect families, in terms of abundance) at each study location/reclamation status combination were plotted to show differences in dominant families. In the case of the reference site at Mesa, only the top two ranked insect families were included because the overall number of insects in the third ranked category was less than 5. The negative binomial regressions and associated bootstrapped confidence intervals were run in Program R [60] using the MASS and ciTools librariers, respectively.

## 3. Results

### 3.1 Vegetation Sampling

Percent cover for each cover class was not statistically different between reclaimed and reference for any class/site combination except for bareground (Riverside and Mesa locations), shrub (Riverside location), and grass (Stewart Point location) as demonstrated by paired t-comparisons with Bonferroni adjustments in Table 1. Grouped bar plots showing the relative percent cover each of cover category/site combination for reference and reclaimed areas are provided in Figure 2.

**Figure 2.**
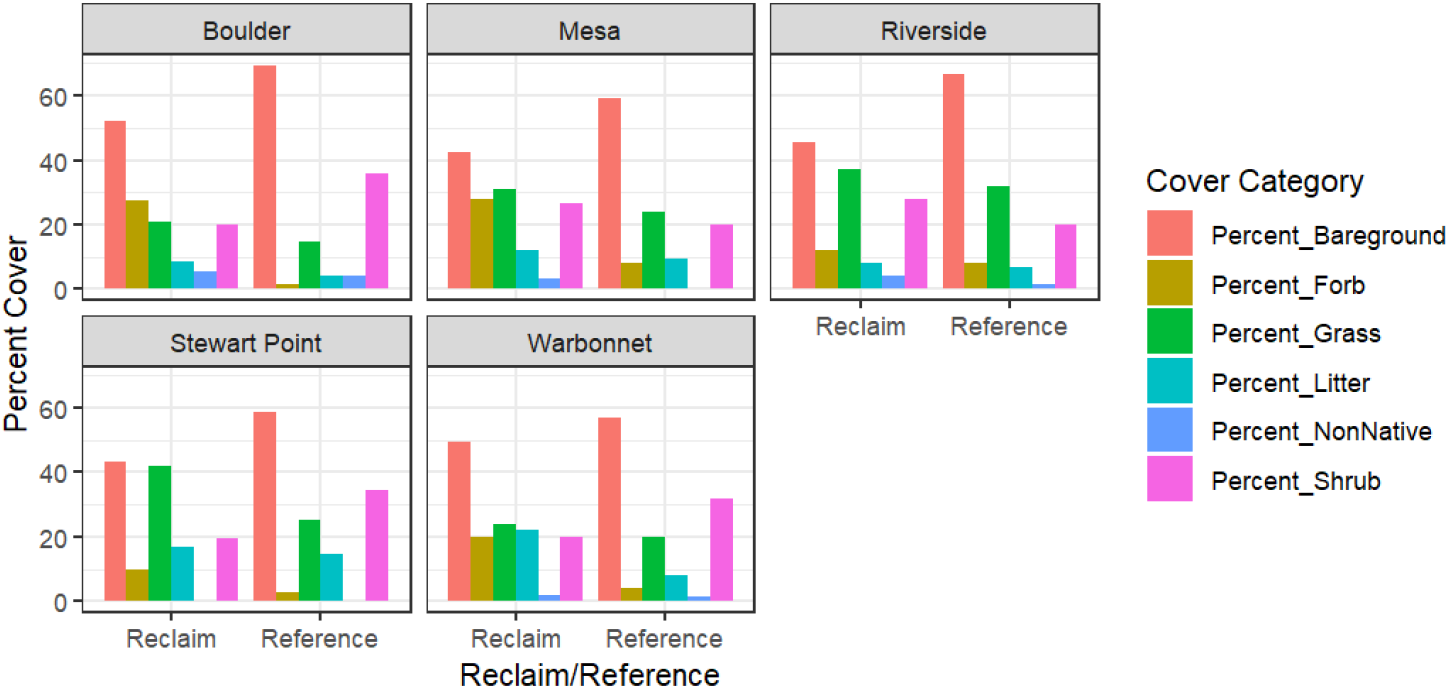
Grouped bar plots showing the relative percent cover each of cover category/site com-bination for reference and reclaimed areas.

### 3.2 Insect Sampling

A total of 2036 individual insects representing 270 species from 71 families across 11 orders were identified across this study. A total of 1557 individuals (76.5%) were found on reclamation sites, whereas 479 (23.5%) were found in reference areas across the entire study. A total of 233 species (86.3% of total) were found on reclamation sites, whereas 121 species (44.8% of total) were found in reference areas across the entire study. A total of 67 families (94.4% of total) were found on reclamation sites, whereas 45 families (63.4% of total) were found in reference areas across the entire study. All 11 orders found in the study were found on reclamation sites, whereas 9 orders were found in reference areas across the entire study.

Insect abundance was significantly greater on reclamation sites than reference sites in three of the five study areas (seeded in 2011, 2013, and 2018) as evidenced by the non-overlapping confidence intervals comparing reclaimed to reference in Figure 3, Panel A. While the reclaimed sites which were seeded in 2012 and 2015 had higher total insect abundance than reference sites, these differences were not statistically important (p>0.05). Both insect family richness and species richness were significantly greater on reclamation sites than reference sites in four of five study areas as evidenced by the non-overlapping confidence intervals in Figure 3, Panels B & C. For both family and species richness, sites seeded in 2012 had higher mean abundance but were not statistically important (p>0.05).

**Figure 3.**
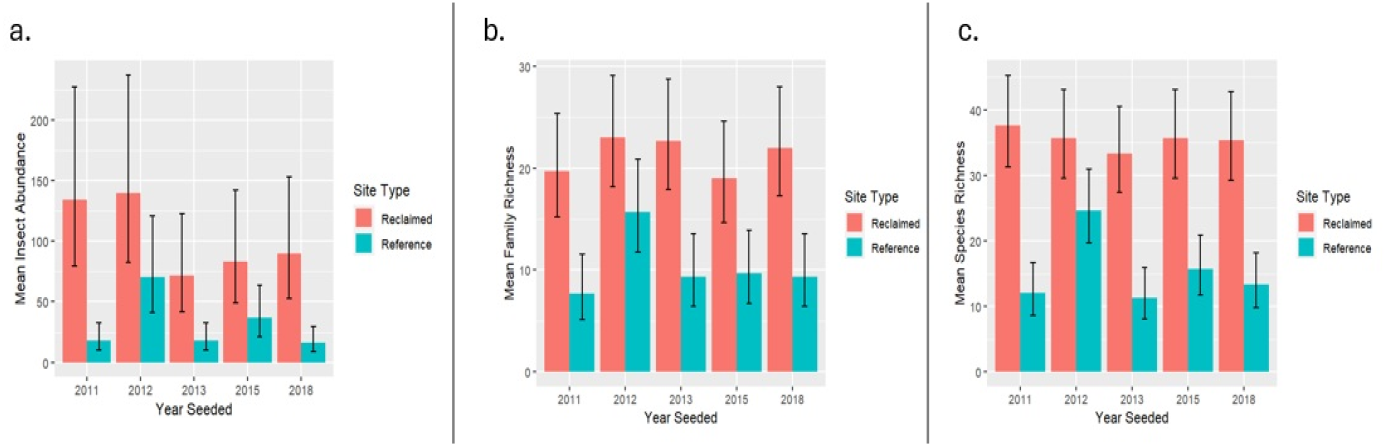
Panel A depicts mean insect abundance in each study area. Panel B depicts mean insect family richness in each study area. Panel C depicts mean insect species richness in each study area.

Dominant insect families varied across management units (Figure 4). In the Boulder management unit (seeded 2011), the dominant families in both the reclamation area and reference sites were *Chalcidoidea, Cicadellidae*, and *Miridae* with all three families found at higher levels in the reclamation area. In the Riverside management unit (seeded 2012), the dominant insect families on the reclamation sites were *Chrysomelidae, Formicidae*, and *Miridae* whereas the dominant insect families in the reference sites were *Miridae, Cicadellidae*, and *Tephritidae*. The Mesa management unit (seeded 2013) saw *Miridae, Tephritidae*, and *Rhyparochromidae* as the dominant insect families on reclaimed sites with *Cicadellidae* and *Miridae* dominant on the reference areas. In the Warbonnet management unit (seeded 2015), the dominant families in both the reclamation area and reference sites were *Chalcidoidea, Cicadellidae*, and *Miridae* with *Cicadellidae* being more abundant in the reference areas and *Chalcidoidea* and *Miridae* more abundant on reclaimed sites. In the Stewart Point management unit (seeded 2018), *Cicadellidae, Formicidae*, and *Miridae* were the dominant insect families on the reclaimed sites while *Cicadellidae, Miridae* and *Chalcidoidea* were dominant in the reference areas – of the common families, more *Cicadellidae* and *Miridae* were found in the reclaimed areas than reference sites.

**Figure 4.**
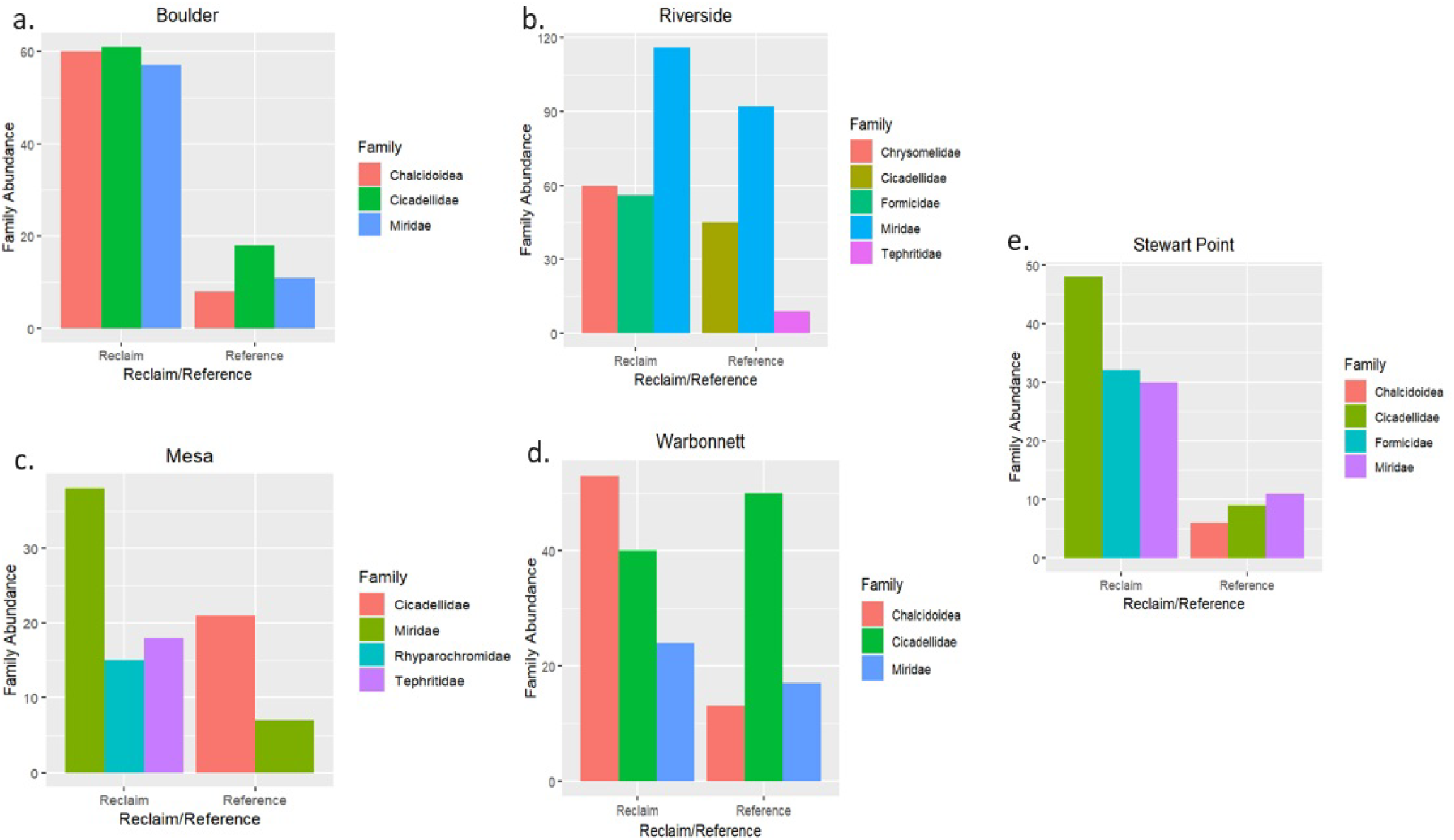
Panel A shows insect family abundance for the only 3 families which made up 62% of the total number of insects in the Boulder (2011 seeded) study area. Panel B shows insect family abundance for the only 5 families which made up >70% of the total number of insects in the Riverside (2012 seeded) study area. Panel C shows insect family abundance for the only 4 families which made up >66% of the total number of insects in the Mesa (2013 seeded) study area. Panel D shows insect family abundance for the only 3 families which made up >63% of the total number of insects in the Warbonnet (2015 seeded) study area. Panel E shows insect family abundance for the only 3 families which made up >65% of the total number of insects in the Stewart Point (2018 seeded) study area.

## 4. Discussion

While little statistically significant differences were found among vegetation cover class groups across this study, it may be important to note that grass, forb, and total vegetation cover had higher mean cover percentages in all study areas. As older stands of sagebrush found in the reference system of the Pinedale Anticline natural gas field have not shown to host large amounts of insect diversity ¬[46, 47], it is perhaps not surprising that insect abundance was significantly greater on well pad groups compared to reference sites in 3 of 5 areas with diversity at the species and family levels being significantly higher on reclaimed sites in 4 of 5 areas studied.

This study was limited to one growing season and conducted only during peak flowering. However, its findings are important as it is the first to examine how reclaimed well pads which are greater than 3-years-old compare to adjacent reference areas in a sagebrush-steppe ecosystem. Previous research in Sublette County natural gas fields clearly showed well pads undergoing reclamation which were 3-years-old and under had more insect abundance and diversity than old stands of reference area sagebrush [46, 47], which is consistent with successional theory suggesting biodiversity peaks in early successional stages [61]. While successional processes develop over long periods of time in cold, arid environments, knowing that reclaimed well pads from 5-12 years old are able to support enhanced biodiversity during peak growing season is encouraging as understanding biodiversity loss and ecosystem restoration have been considered 2 of the top 4 opportunities for future ecological successional studies [62]. The foci of the recent conservation of pattern and process paradigm in rangeland management is on fire and grazing as tools to create patchy mosaics of vegetation communities at the landscape scale [63]. Our findings show promise that biodiversity at the insect level has potential to remain significantly higher on well pads in Sublette County natural gas fields undergoing reclamation for at least 12 years is valuable to the growing body of literature suggesting patchiness in vegetation communities across the landscape is important to rangeland management.

As Greater sage-grouse habitat management and conservation is of utmost importance across Wyoming and the western US [8], our findings are also promising. It is well-documented that insects and forbs play critical roles in the diet of sage-grouse during non-Winter months [15, 64-65]. In addition to insects being important to the sage-grouse diet, it is estimated that 96% of terrestrial birds rear their young solely or primarily on insect protein with declines in insect populations leading to declines in bird populations [66]. As the reclaimed well pads in our study were predominantly comprised of native vegetation, it should be noted that non-native vegetation typically has negative impacts on insects [67]. Therefore, those responsible for reclamation should strive to establish diverse stands of native vegetation.

In addition to our study being limited to only peak growing season during 1 year, our study was also limited in that we used sweep netting to capture insects. It is likely that we would have found different insect communities had we used pitfall traps or other sampling methods [68]. Despite these limitations, our results clearly suggest that natural gas well pads undergoing reclamation within the Pinedale Anticline natural gas field are capable of hosting significantly more insects than reference sites dominated by late successional stands primarily comprised of sagebrush for at least 12 years. Future studies to capture ground-dwelling insects would be beneficial, as would longer-term studies and studies at different points in the growing season.

## 5. Conclusions

As human populations continue to grow, our reliance on natural resources is likely to continue. Ecological reclamation will continue to play an increasingly important role to assist recovery of native biota. As insects provide a plethora of critical ecosystem services, studies to understand how they respond to ecological reclamation efforts will continue to be important. Our results corroborate findings from previous research suggesting that reclamation sites within old successional stands of sagebrush have the ability to support greater insect diversity and abundance, but add to the literature by demonstrating these impacts may last for at least 12 years post initial seeding efforts. Based on our findings, it is recommended that reclamation practitioners in arid and semi-arid sagebrush-steppe ecosystems of western North America continue to use diverse native seed mixes containing forbs and grasses as well as sagebrush.

## Author Contributions

Conceptualization, M.F.C.; methodology, M.F.C., J.A., B.L.R., T.J.R.; reclamation oversight, J.A.; insect identification, A.H.K.; data curation, M.F.C., A.H.K., S.B.; formal analysis, M.F.C., B.L.R., T.J.R.; resources, J.A.; figure creation, T.J.R., B.S.; writing – original draft preparation, M.F.C.; writing – review and editing, M.F.C., J.A., A.H.K., B.L.R., T.J.R., B.M.B., S.B., B.S.; funding acquisition, M.F.C., B.L.R., T.J.R.

## Data Availability Statement

Data will be made available upon reasonable request. Requests should be forwarded directly to M.F.C.

## Acknowledgments

We are grateful to PureWest Energy for providing funding, site access and records related to their reclamation practices. We thank Jesse Dillion (Cedar Creek Associates) for reviewing proposal for funding and discussions about reclamation best practices. We are also grateful to Curt Yanish (Aster Canyon Consulting) and Mike Henn (Sublette County Conservation District) for providing information related to reclamation within the Pinedale Anticline natural gas field. We thank Emily Kress and Timothy Miller for processing, mounting, and labeling insect specimens.

